# Separable functions of Tof1/Timeless in intra-S-checkpoint signalling, replisome stability and DNA topological stress

**DOI:** 10.1101/2019.12.17.877811

**Authors:** Rose Westhorpe, Andrea Keszthelyi, Nicola E. Minchell, David Jones, Jonathan Baxter

**Affiliations:** Genome Damage and Stability Centre, Science Park Road, University of Sussex, Falmer, Brighton, East Sussex BN1 9RQ, U.K

**Keywords:** DNA replication, Replication stress, DNA topological stress, DNA damage

## Abstract

The highly conserved Tof1/Timeless proteins minimise replication stress and promote normal DNA replication. They are required to mediate the DNA replication checkpoint (DRC), the stable pausing of forks at protein fork blocks, the coupling of DNA helicase and polymerase functions during replication stress (RS) and the preferential resolution of DNA topological stress ahead of the fork. Here we demonstrate that the roles of the *Saccharomyces cerevisiae* Timeless protein Tof1 in DRC signalling and resolution of DNA topological stress require distinct N and C terminal regions of the protein, whereas the other functions of Tof1 are closely linked to the stable interaction between Tof1 and its constitutive binding partner Csm3/Tipin. By separating the role of Tof1 in DRC from fork stabilisation and coupling, we show that Tof1 has distinct activities in checkpoint activation and replisome stability to ensure the viable completion of DNA replication following replication stress.

## Introduction

The faithful replication of the genome by the DNA replication machinery is hindered by a range of exogenous and endogenous factors that are capable of disrupting replication forks. Such cellular events are a prominent feature of the early stages of cancer and have been collectively referred to as replication stress (RS) (Halazonetis *et al*, 2008). RS occurs when either the helicase or polymerase activities of the replisome are impeded. Potential impediments include chemical changes to the DNA, stable DNA binding protein complexes, nucleotide deficiency and DNA topological stress (Zeman & Cimprich, 2014; Keszthelyi *et al*, 2016).

On encountering RS, the replisome is thought to be stabilised at the replication fork until the impeding context is removed or bypassed and replication restarted (Marians, 2018). A failure to stabilise the replisome is associated with erroneous processing of the replication fork, leading to either toxic recombination intermediates or disruption of replication restart.

The activation of the checkpoint pathways is essential for replication fork stabilisation and restart (Saldivar *et al*, 2017). Following RS, ATR type kinases (Mec1 in *S. cerevisiae*) are activated by the accumulation of excessive RPA-coated single stranded DNA (Costanzo *et al*, 2003; Zou & Elledge, 2003). ATR/Mec1 then acts with mediator proteins to activate effector kinases that promote fork stability, regulate nucleotide metabolism, and minimise the firing of further replication origins (Pardo *et al*, 2017). The mediator proteins required for effector kinase activation can act either at the fork, commonly referred to as the DNA replication checkpoint (DRC), or independently of the fork, referred to as the DNA damage checkpoint (DDC). Mediator proteins in the DRC include the replisome proteins Mrc1/Claspin (*Sc* Mrc1) (Alcasabas *et al*, 2001) and Tof1/Swi1/Timeless (*Sc* Tof1) (Foss, 2001).

Tof1 appears to act as a nexus for various processes relating to both stabilising the replication fork during RS and ensuring faithful chromosome inheritance. In addition to its role in activating effector kinases, Tof1, along with Mrc1 and the Tof1 interacting protein Csm3/Swi3/Tipin (*Sc* Csm3) are required to stimulate DNA replication *in vivo* and *in vitro* (Tourriere *et al*, 2005; Yeeles *et al*, 2017; Aria *et al*, 2013; Cho *et al*, 2013), and for coupling of helicase and polymerase activities (Katou *et al*, 2003; Bando *et al*, 2009; Errico *et al*, 2007; Smith *et al*, 2009; Lou *et al*, 2008). Independently of Mrc1, the Tof1-Csm3 complex is also required for pausing of replication forks at stable protein-DNA barriers (Tourriere *et al*, 2005; Calzada *et al*, 2005; Hodgson *et al*, 2007) and to focus the action of topoisomerases ahead of the fork to prevent excessive fork rotation during DNA replication (Schalbetter *et al*, 2015). This latter activity has been hypothetically linked to the observation that the C terminus of Tof1 and the type IB topoisomerase Top1 interact (Park & Sternglanz, 1999), potentially ensuring a relative enrichment of Top1 at the replication fork (Schalbetter *et al*, 2015; Keszthelyi *et al*, 2016). Tof1 is also required to recruit Top1 to replicating regions *in vivo* (Shyian *et al*, 2019). In addition, both *tof1Δ* and *csm3Δ* (but not *mrc1Δ*) cells are acutely sensitive to the chemotherapeutic drug camptothecin (CPT) that stabilises the covalently DNA-bound intermediate of Top1(Redon *et al*, 2006; Rapp *et al*, 2010; Hosono *et al*, 2014). The potential linkage between Tof1-Csm3-dependent activities of fork pausing, resolving DNA topological stress and resistance to CPT treatment is unclear.

To attempt to define the molecular relationships between the range of processes that involve Tof1, we have generated a series of truncations across the C terminus of Tof1 and assessed whether these mutations are sufficient for the functions of Tof1 in activating the DRC, replication fork pausing, helicase-polymerase coupling and preventing excessive fork rotation in response to DNA topological stress.

## Results

In order to dissect which domains of the 1238 amino acid (aa) long Tof1 protein are required for each of its characterized functions we generated a series of premature stop codons in a *TOF1* ORF sequence optimised for budding yeast expression (Yeeles *et al*, 2017). Premature stop codons were introduced at aa 627, aa 762, aa 830 aa 997 and aa 1182 (figure 1A). The truncated proteins were expressed from the endogenous TOF1 locus. Truncation of Tof1 at each of these positions generated proteins of the predicted size, which were expressed to levels comparable both with exogenous codon-optimised *wt TOF1* and with endogenous Tof1 protein (figure 1B).

**Figure 1:**
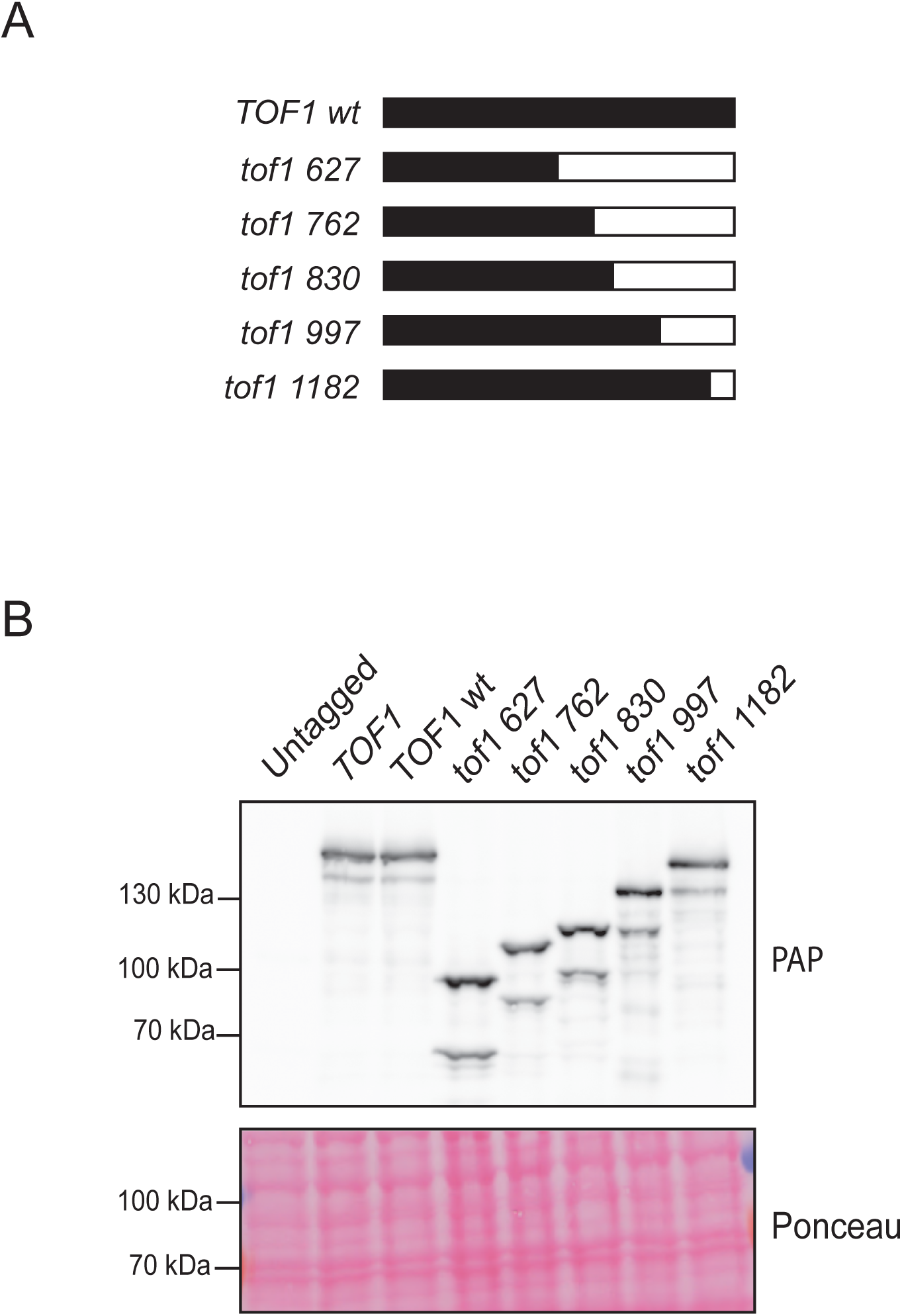
Expression of C-terminal Tof1 truncation mutations. A) Schematic showing length of Tof1 protein expressed by each of the C-terminal truncation mutations compared to wildtype Tof1 (1238 aa). B) Western blot showing expression levels of C-terminal Tof1 truncation mutants. Both endogenous *TOF1* and codon-optimised *TOF1 wt* and *tof1* mutants were C-terminally tagged with TAP epitope and lysates were checked for expression using peroxidase anti-peroxidase (PAP) which immuno-reacts with the protein A portion of TAP tags. Ponceau stain of blotted membrane is shown to illustrate protein content of lanes.

We first set out to establish which of the truncated Tof1 proteins could suppress the excessive fork rotation phenotype of *tof1* deleted cells, as visualised by Southern blotting of replication products (Schalbetter *et al*, 2015). In cells with wildtype function of *TOF1*, episomal plasmids (figure 2A) accumulate only modest levels of DNA catenanes during S phase following Top2 inactivation. This demonstrates that fork rotation is relatively infrequent and therefore that DNA topological stress is primarily resolved by Top1 ahead of the replication fork in this context (Schalbetter *et al*, 2015). In contrast, *tof1Δ* cells accumulate hypercatenated plasmids during S phase indicating that, in this context, DNA topological stress is resolved far more frequently by fork rotation and action of topoisomerases behind the fork (figure 2B). To assess whether the truncated proteins were capable of suppressing hyper-catenation and thus restoring the primacy of Top1 action ahead of the fork, we replaced the *TOF1* gene with each of the *tof1* truncation alleles encoding the truncated proteins into *tof1Δ top2-4* cells containing the plasmid pRS316. Following synchronisation in G1, we cultured the cells at the restrictive temperature to ablate Top2 activity, and released the cells into the cell cycle for one passage through S phase. We then harvested the cells prior to mitosis, preventing further cell cycle progression with the microtubule depolymerising drug nocodazole. We extracted DNA and assessed the frequency of DNA catenation introduced into the plasmid using two-dimensional gel electrophoresis and Southern blotting. As expected, expression of *TOF1 wt* completely suppressed the excessive fork rotation and DNA catenation of *tof1Δ top2-4* cells (Figure 2C). Expression of *tof1 1182* which lacks only the final 57 amino acids of the C terminus of Tof1 also fully rescued the excessive fork rotation (Figure 2H). However, expression of Tof1 proteins truncated at 627, 762, 830 and 997 did not rescue this effect (Figure 2D-G). This indicates that the region of Tof1 between aa 997 and aa 1182 is required to suppress excessive fork rotation. This matches the region of Tof1 (997-1226) that is capable of supporting a two-hybrid interaction with Top1 (Park & Sternglanz, 1999) and recruiting Top1 to the replication fork (Shyian *et al*, 2019). This is consistent with the model that preferential recruitment of Top1 to the replisome stimulates its action ahead of the fork (Schalbetter *et al*, 2015; Keszthelyi *et al*, 2016).

**Figure 2:**
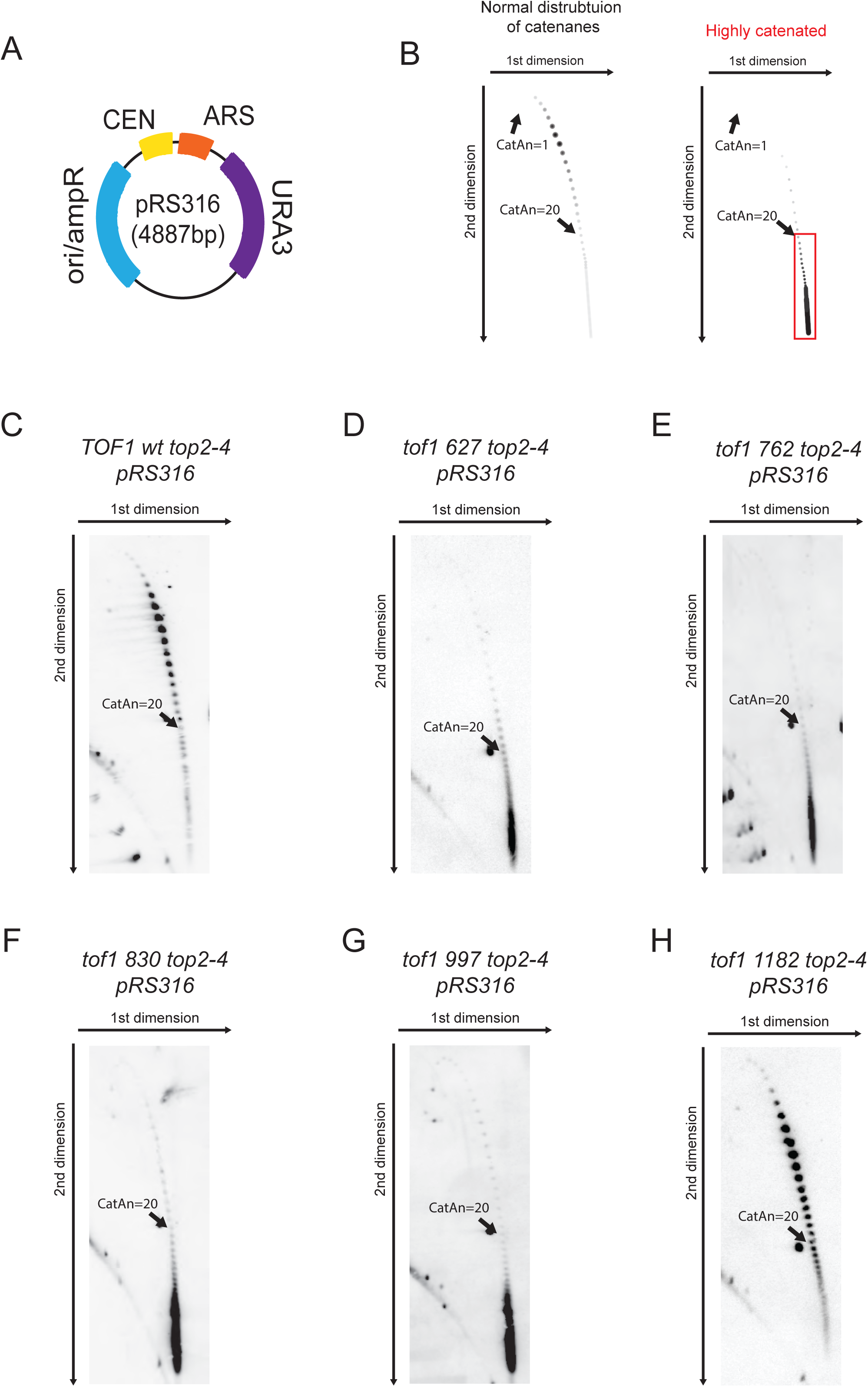
Suppression of hyper-catenation on episomal plasmids in Top2-inactivated cells requires the far C terminal region of Tof1. A) Plasmid pRS316 used for DNA catenation analysis. B) Schematic showing the representative increase in DNA catenation of plasmid pRS316 in *tof1Δ top2-4* cells compared to *top2-4* cells as visualized by Southern blotting. Region of hypercatenated plasmids highlighted in red box. C-H) DNA catenation analysis of pRS316 in *top2-4* cells expressing C) *TOF1 wt* D) *tof1 627* E) *tof1 762* F) *tof1 830* G) *tof1 997* H) *tof1 1182*

To further examine the functionality of the Tof1 truncated proteins we next set out to determine which of the mutants were able to support replication fork pausing at a replication block. We cloned the *S. cerevisiae* rDNA region that contains the replication fork barrier (RFB) (corresponding to Chromosome XII sequence : 459799 – 460920) into the multicopy yeast episomal plasmid pRS426 (figure 3A). Transplanting the RFB sequence into an artificial location pauses replication forks in a Fob1 dependent manner, but does not fully arrest ongoing replication (Calzada *et al*, 2005). It therefore appears to act in a similar manner to other endogenous protein pausing sites (Hodgson *et al*, 2007)(cartoon example shown in figure 3B). Cells without Tof1 did not pause at the RFB site on the plasmid (figure 3C), whereas cells expressing *TOF1 wt* fully supported pausing at this location (figure 3D). Expression of *tof1 830, tof1 997 and tof1 1182* also fully supported pausing (figure 3G-I).

**Figure 3:**
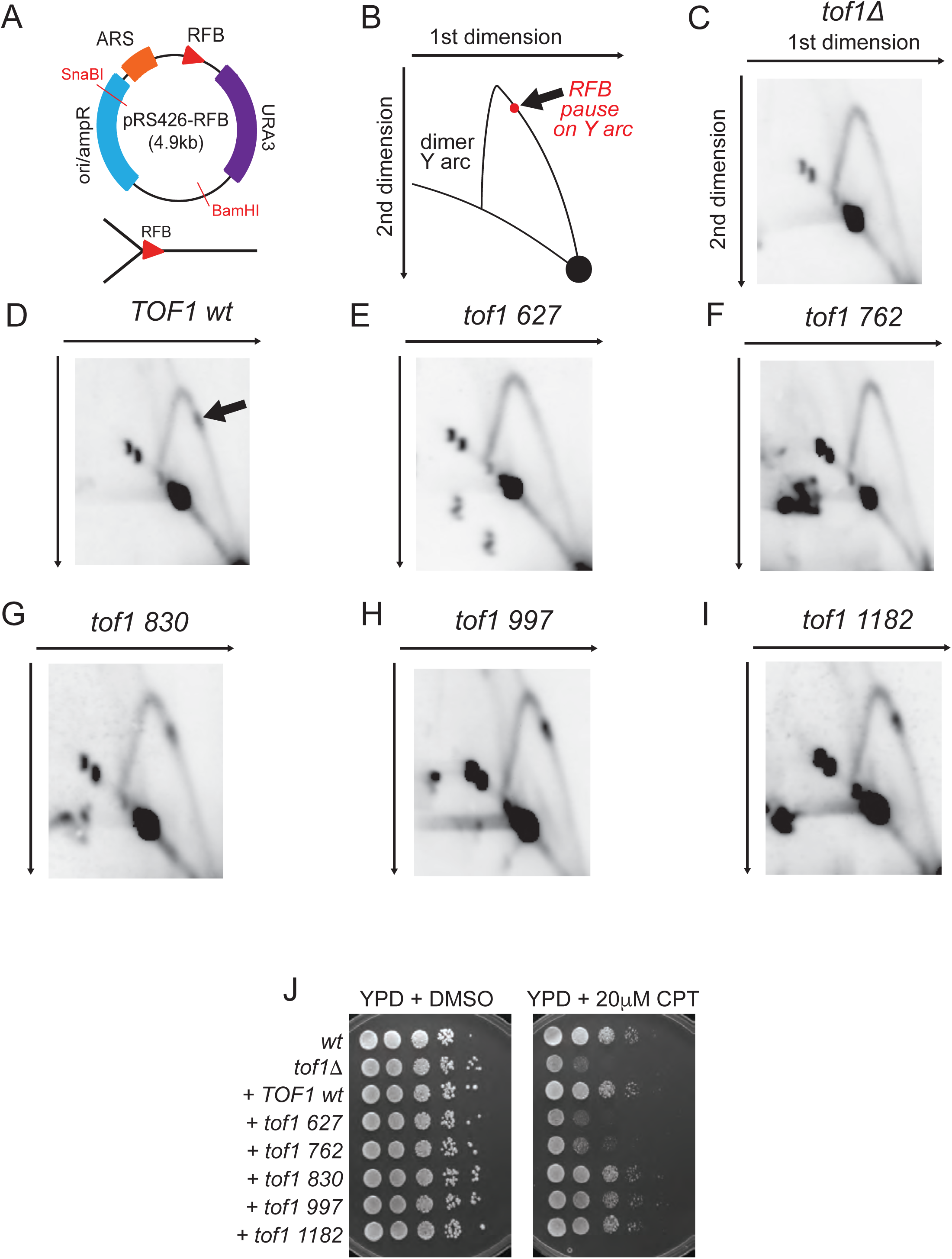
Recovery of fork pausing and resistance to camptothecin requires the central region of Tof1 but not the far C terminal region. A) Plasmid pRS426-RFB used for analysis of fork pausing at the RFB. The two unique restriction digest sites used to linearize the plasmid for 2D replication fork gel analysis and the Y structure anticipated to be generated by fork pausing at the RFB pause site are shown. B) Schematic of the replication fork Y arc predicted to be generated by pRS426-RFB digestion with BamHI and SnaBI in wildtype cells and resolved by 2D gel electrophoresis. Arrow indicates the accumulation of replication intermediates generated by pausing at the RFB on pRS426. C-I) 2D gel analysis of pRS426-RFB in exponentially growing cells containing C) *tof1Δ* D) *TOF1 wt* E) *tof1 627* F) *tof1 762* G) *tof1 830* H) *tof1 997* I) *tof1 1182* J) Viability spot assay of wildtype (*wt*), *tof1Δ, TOF1 wt, tof1 627, tof1 762, tof1 830*, *tof1 997*, and *tof1 1182* cells grown on YPD + DMSO or YPD plus 20 µg/ml camptothecin plates for 48 hours at 25°C.

However, expression of *tof1 627* and *tof1 762* did not rescue pausing at the RFB site (figures 3E and 3F). Together this data indicates that the last 408 aa of Tof1 are not required for replication fork pausing at protein barriers to DNA replication and therefore that the role of Tof1 in replication fork pausing is independent of its role in inhibiting fork rotation.

Deletion of *TOF1* results in hypersensitivity to CPT, an agent which stabilises the Topoisomerase 1 covalent complex (Top1cc) to DNA, forming a DNA protein cross-link (DPC). Deletion of *CSM3* results in similar sensitivity to CPT treatment while *mrc1*Δ cells display only mild sensitivity to this agent (Redon *et al*, 2006). Therefore, CPT sensitivity appears to be a marker for functions of the Tof1-Csm3 heterodimer that are independent of their association with Mrc1. The cause of the acute toxicity of *tof1Δ* or *csm3Δ* cells to CPT is currently unclear. On the leading strand a DPC and an adjacent single strand DNA break would be predicted to either inhibit replication fork progression or generate a double strand break (Strumberg *et al*, 2000; Sparks *et al*, 2019). Alternatively, CPT treatment has also been found to cause increased DNA topological stress on cellular DNA (Koster *et al*, 2007), potentially generating a situation where efficient recruitment of topoisomerases to the fork by Tof1 is required for ongoing elongation. Our data has established that *tof1* truncations that cannot support relaxation of DNA topological stress through Top1 recruitment can still support fork pausing at a RFB complex. We therefore examined how expression of each of the truncation mutations suppressed the sensitivity of *tof1Δ* cells to CPT. Cells expressing *tof1 627* or *tof1 762* were highly sensitive to CPT while expression of *tof1 830, tof1 997* and *1182* provided *wt* levels of resistance to CPT (figure 3J). Therefore, truncation mutants that do not support Tof1’s role in promoting Top1 action ahead of the fork (*tof1 830* and *tof1 997*) are normally resistant to CPT. Conversely, *tof1* truncations that do not support replication fork pausing (*tof1 627* and *tof1 762*) cannot suppress CPT sensitivity. We conclude that CPT sensitivity is closely associated with the ability of the Tof1-Csm3 complex to support pausing of the fork at a protein block to replication.

Replisome pausing induced by nucleotide depletion with Hydroxyurea (HU) also requires Tof1 function. Following acute treatment of cells with HU both Tof1 and Mrc1 are required to maintain the coupling of helicase and polymerase activities at the fork (Katou *et al*, 2003). In the absence of Tof1 or Mrc1 the helicase advances beyond the point of nascent DNA incorporation, indicating uncoupling of helicase and polymerase activities in the replisome (Katou *et al*, 2003). This uncoupling is predicted to lead to increased binding of the single stranded DNA binding protein, RPA1, to the single stranded regions generated by uncoupled helicase action. We have experimentally confirmed this prediction using RPA1 ChIP-SEQ in HU treated wildtype, *tof1* and *mrc1* cells. Release of cells arrested in G1 with alpha factor into media containing 200 mM HU led to strongly increased RPA1 chromatin binding around replication origins in *tof1*Δ and *mrc1*Δ cells compared to wildtype (figure 4A). Expression of *TOF1 wt* and *tof1 830* both suppressed the accumulation of RPA1 around early firing origins (figure 4B). Thus, expression of the *tof1 830* truncation mutant was sufficient to ensure helicase and polymerase coupling. However, expression of either *tof1 762* or *tof1 627* still led to increased RPA1 around early origins, indicating helicase-polymerase uncoupling. Notably the elevated level of RPA1 around origins in either *tof1 762* or *tof1 627* was consistently less than the level of RPA1 observed in *tof1*Δ cells (figure 4B), indicating that they retained some function in coupling of helicase and polymerase activities, potentially through their mediation activity in the DRC (see below).

**Figure 4:**
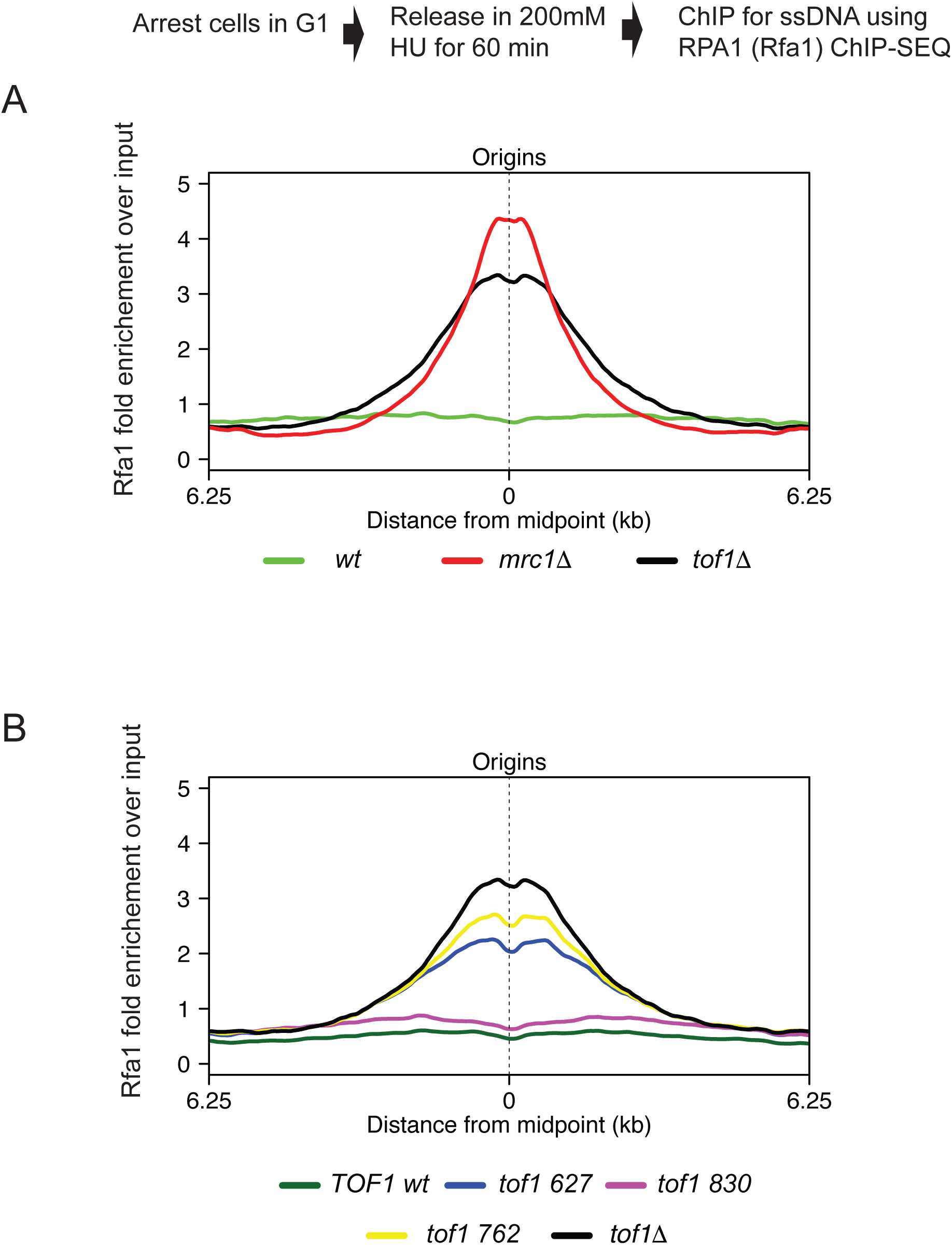
Coupling of helicase and polymerase activities requires the central region of Tof1 but not the far C terminal region. To examine the extent of fork uncoupling we assayed ChIP of RPA1 (*Sc*Rfa1) following 200 mM hydroxyurea (HU) arrest of cell proliferation A) Schematic of experimental set up and meta-analysis of enrichment of RPA1 (*Sc*Rfa1) around origins in *wt, tof1Δ* and *mrc1Δ* cells following release of G1 cells into media containing 200 mM HU for 60 min. B) Meta-analysis of enrichment of RPA1 (*Sc*Rfa1) around origins in *TOF1 wt, tof1Δ tof1 627, tof1 762* and *tof1 830* cells following release of G1 cells into media containing 200 mM HU for 60 min.

We next set out to establish whether expression of the truncated proteins was sufficient to support the mediator function of Tof1 in activation of the DRC. In *S. cerevisiae,* sustained activation of the effector kinase Rad53 in S phase following HU treatment can be through the DRC mediated by Mrc1 and Tof1, or through the Rad9-mediated DDC (Pardo *et al*, 2017). Therefore, loss of detectable activation of Rad53 in response to HU only occurs in cells lacking both *tof1* and *rad9* function. Conversely, HU-dependent Rad53 activation can be rescued by the activity of either Tof1 or Rad9 protein in *tof1Δ rad9Δ* cells. We assayed whether expression of any of the *tof1* truncation mutants could rescue Rad53 activation in response to HU treatment in *tof1Δ rad9Δ* cells, using the Rad53 phospho-mobility assay (Pellicioli *et al*, 1999). In these experiments, treatment of exponentially growing *wt* cells with 200 mM HU led to a robust auto-phosphorylation-linked mobility shift of Rad53 protein as visualised by western blot. This shift was partially attenuated in *tof1Δ* and completely lost in *tof1Δ rad9Δ* cells (figure 5A). Interestingly, expression of all of the truncation mutants rescued the phosphorylation-linked mobility shift of Rad53 in response to HU (figure 5A).

**Figure 5:**
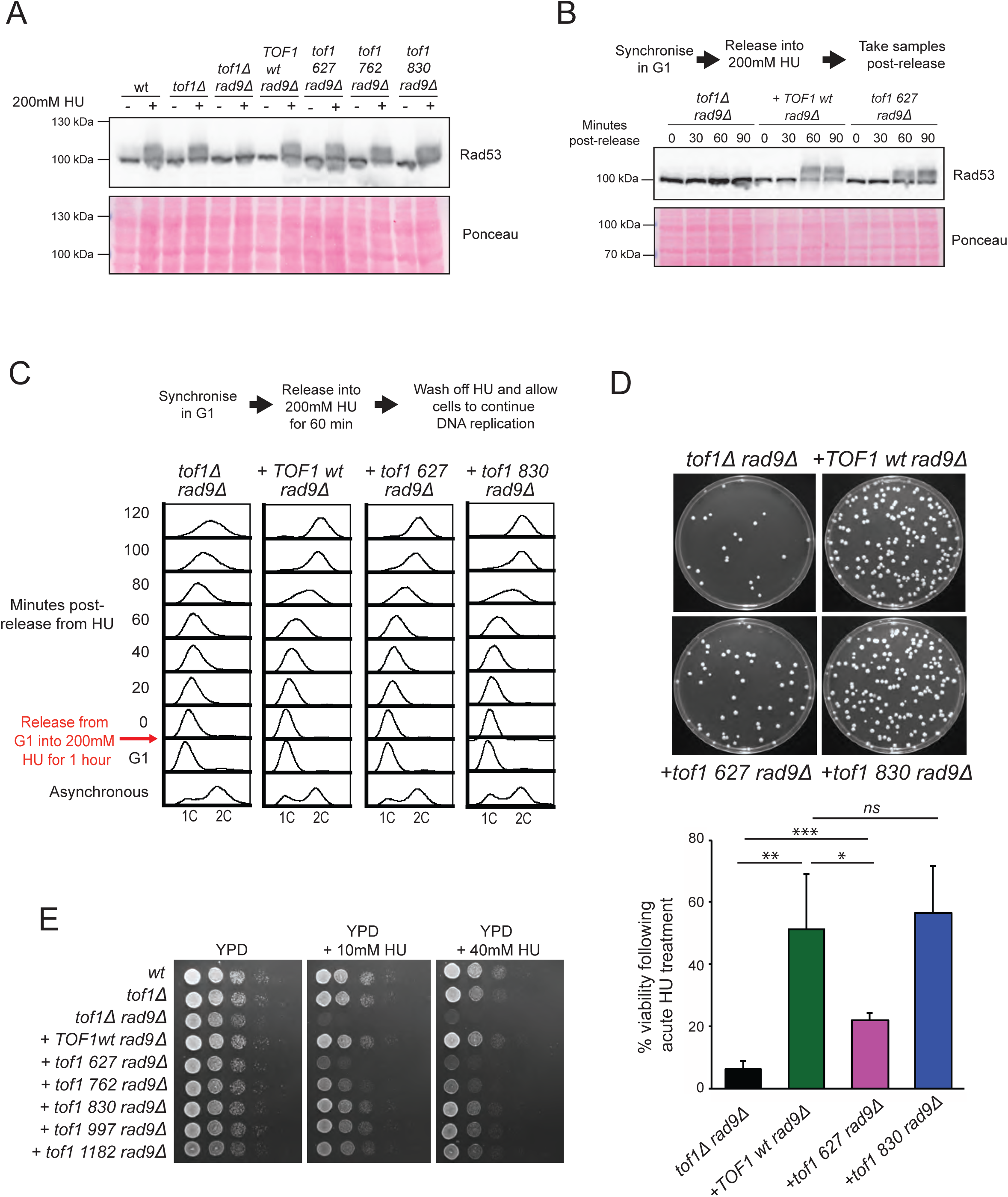
Following HU arrest the N terminal half of Tof1 is sufficient for DRC activation and resumption of DNA synthesis but not cell viability in DDC deficient (*rad9Δ*) cells. A) Rad53 activation assay following HU treatment. Activation of Rad53 causes autophosphorylation that results in reduced mobility of the Rad53 protein. The relative mobility of Rad53 in each of the indicated cell types either in exponentially growing cells or following a 2-hour treatment with 200 mM HU was assayed by western blot using anti-Rad53 antibodies (top). Ponceau stained membrane of western blot shown in top panel to indicate relative protein levels (bottom). B) (Top) Time course of Rad53 activation following release from G1 into media containing 200 mM HU. (Bottom) Ponceau stained membrane of western blot shown in top panel to indicate relative protein levels. C) Recovery of DNA synthesis in cells following cell cycle arrest induced by 200 mM HU. The indicated cell mutants were arrested in 200 mM HU following release of a G1 synchronised culture. After 60 minutes in HU, the HU containing media was washed off and cells allowed to recover and resume passage through S phase. Shown are FACS profiles of DNA content before and after the HU arrest point. Time points indicate the time in minutes after release from the HU arrest (T0 indicates time from the first wash). D) Viability of cells following cell cycle arrest induced by acute HU treatment. G1-arrested cells were released from the block into YPD media containing 200 mM HU for 1 hour, after which the cultures were diluted and plated to YPD plates. Colony number was counted 48 hours after plating and results from 4 repeats were quantified and displayed in histogram (bottom). P values: *=<0.05, **=<0.01, ***=<0.001. E) Viability of cells during chronic treatment with HU. Viability spot assay of *TOF1 wt*, *tof1Δ, tof1 627, tof1 762, tof1 830*, *tof1 997* and *tof1 1182* cells on YPD or YPD plus 10 or 40 mM HU for three days at 25°C.

Therefore, the first 627 aa of Tof1 retains the mediator function of the protein in the DRC. Previous studies have shown that hypo-morphic alleles of some checkpoint components delay but do not prevent the full activation of Rad53 following treatment with HU (Hustedt *et al*, 2015). To assess whether this was the case for the *tof1 627* allele we assayed protein extracts from cells taken at sequential time points following release into 200 mM HU for Rad53 activation (figure 5B). We did not observe a delay in activation of Rad53 in the *tof1 627* allele compared to wildtype Tof1, consistent with this allele activating the intra-S-checkpoint with wildtype kinetics.

The separation of function observed between the roles of Tof1 in checkpoint signalling and in replisome coupling/pausing provided us with an opportunity to differentiate between the contribution of these two Tof1 functions to genome stability following RS induced replisome stalling. Following treatment of budding yeast with 200 mM HU, replication forks stall shortly after origin firing, activating the DRC. DRC activation is essential to maintain cellular viability during incubation in HU (Lopes *et al*, 2001; Tercero *et al*, 2003). Checkpoint-dependent viability is thought to be due to regulation of several pathways including inhibition of late origin firing (Paciotti *et al*, 2001), inhibition of nucleases that could process the arrested fork (Segurado & Diffley, 2008; Cotta-Ramusino *et al*, 2005) and stabilisation of replisome components at the arrested fork (Lopes *et al*, 2001; Tercero *et al*, 2003). The role of Tof1 in maintaining genome stability in response to RS could be through checkpoint signalling, through a general role in maintaining replisome integrity, or a combination of the two. To investigate the contribution of these two roles to genome stability we compared the response to arrest and subsequent wash off of 200 mM HU in *tof1*Δ *rad9*Δ cells, where the checkpoint activation of Rad53 is ablated, to *tof1*Δ *rad9*Δ cells complemented with either *TOF1 wt*, t*of1 627* or *tof1 830*, all of which generate Tof1 protein capable of activating the DRC in *tof1*Δ *rad*Δ*9* cells (figure 5C and 5D). Using FACS analysis of DNA content, we observed that uncomplemented *tof1*Δ *rad9*Δ cells were unable to complete DNA replication following removal of 200 mM HU, consistent with previous analysis of checkpoint-defective cells (Desany *et al*, 1998; Lopes *et al*, 2001) (figure 5C). Complementation of these cells with wildtype Tof1 resulted in rapid completion of DNA replication (figure 5C). In addition, complementation of *tof1*Δ *rad9*Δ cells with either of the checkpoint competent truncation mutants *tof1 627* or *tof1 830* also resulted in rapid apparent completion of DNA replication (figure 5C), consistent with the checkpoint function of Tof1 alone being sufficient for resumption of DNA replication following HU arrest. In addition to testing DNA content we also re-plated these cells following release from the acute HU treatment onto YPD plates to assess whether they could survive the treatment. As expected, *tof1*Δ *rad9*Δ cells failed to form colonies following acute HU treatment whereas *TOF1 wt rad9Δ* cells displayed efficient colony-forming capability (figure 5D). Despite their apparent resumption in DNA replication following acute HU treatment, *tof1 627* cells generally failed to recover from the HU-induced arrest, although they were more viable than *tof1*Δ *rad9*Δ cells (figure 5D). This indicates that despite DRC activation and apparent efficient resumption of DNA replication, *tof1 627* expressing cells were still subject to frequent unrecoverable lesions following fork stalling. Consistent with this interpretation both *tof1 627* and *tof1 762* cells were acutely sensitive to chronic treatment with HU compared to the other *tof1* truncation mutants (figure 5E).

FACS analysis of DNA content cannot distinguish if any increases in DNA content are due to restart of arrested replication forks or due to new origins firing. To investigate this issue, we carried out sync-seq analysis (Müller *et al*, 2014) (Conrad Nieduszynski personal communication) to compare copy number differences across the genome between cells arrested in 200 mM HU and cells 80 minutes following wash off of HU, recovery and resumption of DNA replication (figure 6A). Analysis of *TOF1 wt rad9Δ*, *tof1 627 rad9Δ*, *tof1 830 rad9Δ* and *tof1Δ rad9Δ* cells arrested in HU all showed similar copy number in the immediate vicinity of early firing origins as expected (figure 6B left). Further sync-seq analysis of the same culture 80 minutes after HU removal showed that *TOF1 wt* and *tof1 830* cells had completed replication around origins fired before HU arrest as anticipated, indicating that stalled forks in these cells were able to restart replication following release from the HU block (figure 6B right). As predicted, *tof1Δ rad9Δ* cells had a much smaller increase in copy number around origins consistent with poor levels of recovery or restart from HU-arrested forks in this genetic background (Figure 6B right). In contrast to these two extremes, *tof1 627* cells exhibited a similar pattern of increased copy number around origins fired in HU to *wt* and *tof1 830,* but often failed to increase copy number to the same level as seen in late replicating regions (figure 6B bottom right).

**Figure 6:**
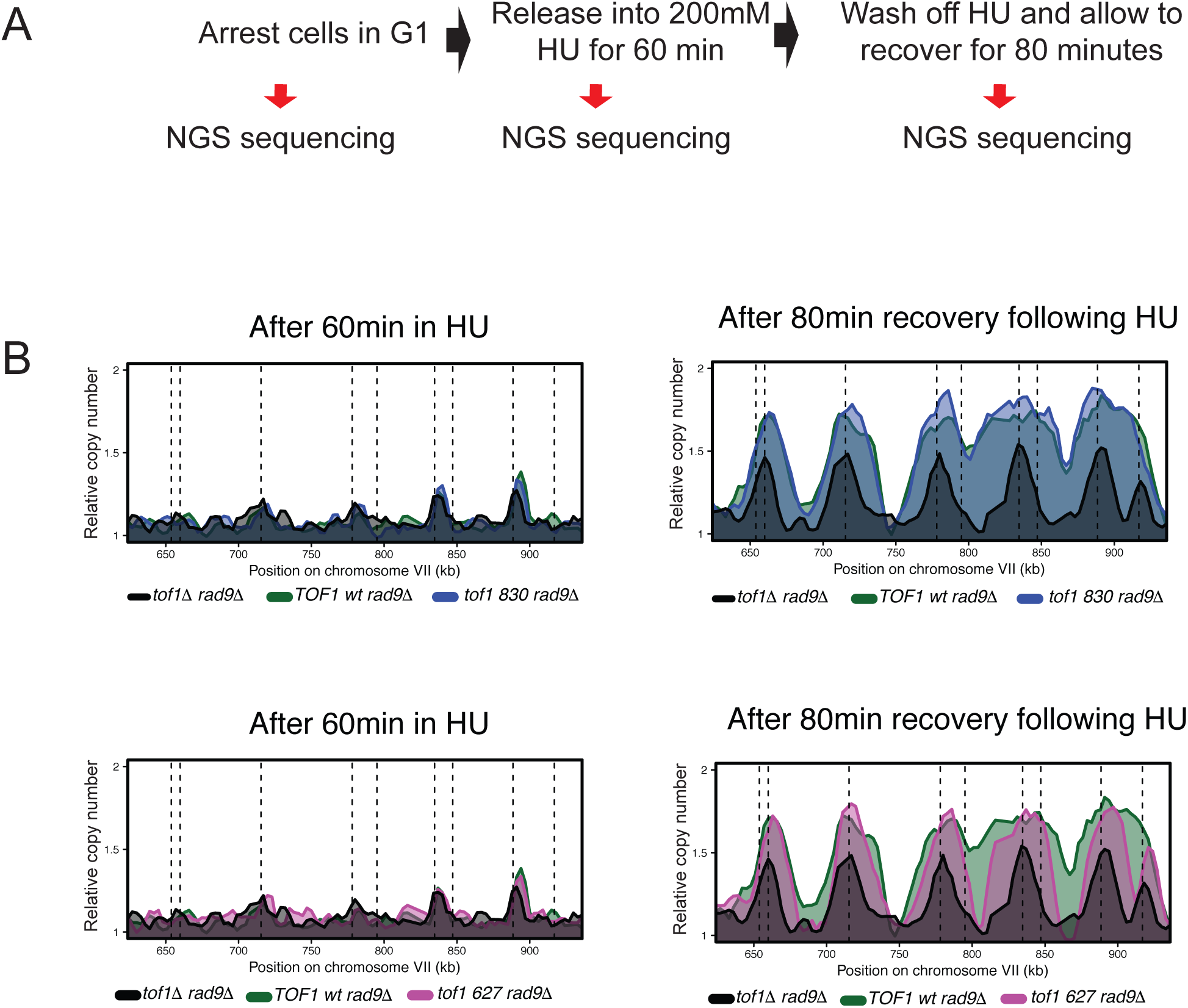
Replication in *tof1 627* cells is reduced after release from HU treatment in *rad9Δ background*. A) Experimental set up for sync-seq experiments in cells arrested in 200 mM HU and released to follow replication fork restart. B) Overlaid relative copy number of *TOF1 wt rad9Δ* and *tof1Δ rad9Δ* compared to *tof1 830 rad9Δ* (top panels) and *tof1 627* (lower panels) after 60 min exposure to 200mM HU (left panels) and 80 min after wash off of HU allowing the cells to recover and resume DNA replication (right panels). Dashed lines represent known origins.

These findings indicate that Tof1 proteins that can mediate DRC activation can promote efficient restart around HU-stalled forks. However, replication forks associated with Tof1 proteins that cannot support replisome coupling or pausing then go on to exhibit elongation defects that often prevent the completion of DNA replication. This leads to toxic lesions if the Rad9-dependent repair pathways are absent.

In summary, we have found that the function of Tof1 in checkpoint activation only requires the N-terminal half of the Tof1 protein. In contrast the roles of Tof1 in supporting fork pausing, suppressing CPT sensitivity, coupling of helicase and polymerase activities and maintaining fork stability following restart all require amino acids 627-830 of Tof1. Interestingly, the role of Tof1 in resolving DNA topological stress ahead of the fork is confined to the region beyond aa 997. Amino acids 1-813 of human Timeless are the minimal region required for interaction with the human homologue of Csm3, Tipin (Holzer *et al*, 2017). Therefore, loss of amino acids 627-830 of Tof1 could partially destabilise the interaction between Tof1 and Csm3. We assessed the stability of the Tof1-Csm3 interaction by immunoprecipitation of TAP-tagged Tof1 truncation proteins and western blotting for Csm3 (figure 7). We observed robust interaction between Tof1 and Csm3 in *TOF1 wt*, *tof1 830*, *tof1 997* and *tof1 1182* expressing cells. In contrast we were unable to detect a stable interaction between Tof1 and Csm3 in *tof1 627* and *tof1 762* cells. Previous studies have shown that loss of Tof1/Timeless family proteins leads to a decrease in the detectable level of Csm3/Tipin protein suggesting that the interaction between the two is required for Csm3/Tipin stability in cells (Chou & Elledge, 2006; Bando *et al*, 2009). To assess for loss of stable Csm3 protein, we western blotted for Csm3 in input extracts of cells expressing WT and truncated Tof1 proteins (figure 7). Expression of *TOF1 wt, tof1 830, tof1 997* and *tof 1182* all supported normal levels of Csm3 protein. Notably expression of *tof1 627* and *tof1 762* led to partially reduced levels of Csm3 in cells (figure 7). This is consistent with the *tof1 627* and *tof1 762* truncations compromising the stability of the interaction between Tof1 and Csm3, leading to partial destabilisation of the Csm3 protein.

**Figure 7:**
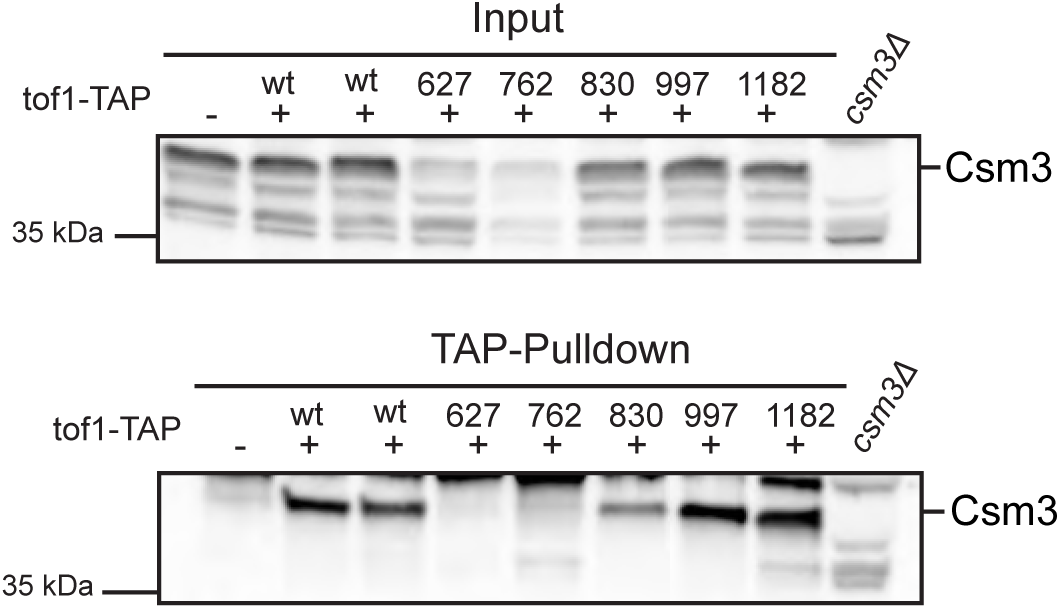
Amino acids 627-830 of Tof1 are required for its stable association with Csm3. Pulldown of Csm3 in TAP-tagged Tof1 truncation mutants. Lysates from cells containing C-terminally TAP-tagged *tof1* truncation proteins were immunoprecipitated and the resulting eluates were western blotted using antibodies to Csm3.

## Discussion

Defining how the Timeless protein functions to maintain genome stability, both generally and following RS has been complicated by its pleiotropic functions. Here we have shown that distinct regions of Tof1 are required for its roles during DNA replication.

We find that the far C terminal region (>997aa) of Tof1 is required to supress excessive fork rotation in cells. This region is also sufficient to mediate a two-hybrid interaction with the type 1B topoisomerase Top1 (Park & Sternglanz, 1999). During DNA replication Top1 can only act ahead of the replication fork to facilitate unwinding of the parental DNA duplex (Schalbetter *et al*, 2015; Keszthelyi *et al*, 2016). Therefore, the recruitment of Top1 to the replisome via the C terminal region of Tof1 will bias topoisomerase action to the region ahead of the fork, providing a ready explanation for why this region is required to suppress the alternate pathway of duplex DNA unwinding, specifically fork rotation and the action of Type II topoisomerase behind the fork. During preparation of this study others have also shown that a similar C terminal region of Tof1 is required to recruit Top1 to active replication forks, consistent with this model (Shyian *et al*, 2019). Interestingly, the SV40 T antigen helicase also directly recruits Top1 to the SV40 replisome to facilitate DNA unwinding (Simmons *et al*, 1996) suggesting that direct replisome recruitment of Top1 could be a general feature of eukaryotic replisomes. It remains to be seen if topoisomerase recruitment to the eukaryotic replisome is normally achieved via Timeless family proteins or if this function could be swapped between different replisome components in different organisms.

Distinct from the far C terminus, the middle region of Tof1 between amino acids 627 and 830 are required for fork pausing at the RFB protein barrier and also to maintain coupling of helicase and polymerase activities during HU induced RS. Expression of *tof1 627* and *tof1 762* truncations ablates the interaction between Tof1 and Csm3. This also appears to be linked to a general partial loss of Csm3 protein levels in cells, suggesting that at least part of the function of this region is to stabilise the Tof1-Csm3 interaction. We speculate that the Tof1-Csm3 heterodimer plays a central role in replisome structure that facilitates coordination of CMG and replicative polymerases consistent with observations across several systems (Katou *et al*, 2003; Cho *et al*, 2013; Errico *et al*, 2007). Current data argues that functional consequences of the Tof1-Csm3 interaction extend beyond the mapped 627-830 aa region of Tof1. For example, we have previously shown that *csm3Δ* cells do not preferentially target Top1 activity to the region ahead of the fork, despite full length Tof1 still being expressed (Schalbetter *et al*, 2015). This argues that the Tof1-Csm3 interaction is required to appropriately configure the Tof1-Top1 interaction within the replisome. This region of Tof1 that stabilises its interaction with Csm3 is also required to suppress the sensitivity of *tof1Δ* cells to the Top1cc complexes generated by CPT. Since the C-terminal region required for the Tof1-Top1 interaction is dispensable for suppression of CPT sensitivity in *tof1Δ* cells, it appears that the coordination of Top1 action with the replisome by Tof1 does not influence CPT sensitivity. Rather our data argue that CPT sensitivity is closely linked with stable fork pausing and helicase-polymerase coupling. We speculate that Tof1 activity allows the fork to pause when encountering a Top1cc complex on the leading strand, preventing polymerase runoff (Strumberg *et al*, 2000) and the generation of a potentially lethal single ended DNA DSB.

Our data argue that the roles of Tof1 in responding to DNA topological stress are distinct from its roles in pausing at replication fork blocks. In contrast to our data, others have recently shown a partial loss of pausing at the endogenous RFB through expression of a Tof1 protein truncated at 981 amino acids, which cannot recruit Top1 to the fork. At present the reason for the variation in fork pausing efficiency of our truncations at 830 and 997 relative to a truncation at 981 is unclear. Further additional loss of pausing at the endogenous RFB was observed in the absence of Top2 and the presence of the truncated Tof1 protein (Shyian *et al*, 2019). This suggests that, at least at the endogenous RFB, the action of either Top1 or Top2 ahead of the fork promotes fork pausing. Whether this model reflects a general model for fork pausing at protein blocks to replication or is specific to the endogenous RFB remains to be determined.

Finally, we find that the N-terminal half (627 aa) of Tof1 is sufficient for efficient intra-S-checkpoint signalling in response to HU. Both Mrc1 and Tof1 have been identified as required for efficient checkpoint signalling in HU (Alcasabas *et al*, 2001; Foss, 2001). Mrc1 alone appears capable of facilitating the sensor/effector kinase interaction (Berens & Toczyski, 2012), whereas evidence that Tof1/Timeless acts as a mediator of the checkpoint in the absence of Mrc1/Claspin is lacking. Rather it appears more likely that Tof1 is required to promote association of Mrc1 with the replisome (Bando *et al*, 2009; Yeeles *et al*, 2017), thus indirectly promoting the role of Mrc1 in the DRC. We speculate therefore that the N-terminal half of Tof1 is sufficient for Mrc1 to be associated with the replisome in a manner compatible with mediating DRC signalling.

DRC activity following fork stalling ensures that the replisome and fork structure is maintained in a form capable of efficiently restarting. Tof1’s role as a mediator protein provides a straightforward explanation of why its absence results in problems in fork restart (Tourriere *et al*, 2005). However, the replisome instability caused by loss of Tof1 also suggests that replisome structure without Tof1 is not proficient in restart following HU arrest irrespective of the whether or not the checkpoint is activated. The separation of function shown by the *tof1 627* mutant, which is proficient for checkpoint signalling but not proficient for replisome coupling or fork pausing, has allowed us to show that Tof1 has important roles in completing DNA replication following RS beyond its role in DRC activation activity and DRC dependent restart. Our data argues that Tof1 is required to stabilise restarted forks for subsequent elongation of the replisome. The reason for fork failure is unclear but we speculate that *tof1* mutants deficient in coupling helicase and polymerase activity in HU are also defective in efficient recoupling during recovery from HU treatment, leading to frequent fork failure distal from the restart site. Notably, the sensitivity of *tof1* mutants to HU is highly dependent on the activity of Rad9, consistent with the Rad9 dependent repair pathways being required for *tof1* linked DNA lesions. Although our data show that Tof1 has roles in recovering from RS beyond mediation of the DRC, we cannot rule out a role for Tof1 downstream of DRC activation. The phosphorylation state of Tof1 is partially dependent on Mec1/Tel1 activity (de Oliveira *et al*, 2015) and it is known that Tof1 phosphorylation can influence fork pausing (Bastia *et al*, 2016). Therefore, checkpoint dependent phosphorylation of the C terminal half of Tof1 could be important for recovery of the fork following RS.

In summary, our data argues that the N-terminal half of Tof1 is linked to Mrc1 replisome functions, and that the far C terminus regulates Top1 association with the replisome. We find that an internal region of Tof1, between aa 627 and aa 830 that is required for stable binding to Csm3, is crucial for fork pausing and replisome coupling of helicase and polymerase activities. These data indicate that Tof1 has roles both in co-ordinating general replisome architecture independently through both Mrc1 and Csm3 interactions, as well as specifically recruiting other activities such as Top1 to the replisome.

## Abbreviations

aa: amino acids
CPT: Camptothecin
DDC: DNA damage checkpoint
DPC: DNA protein cross-link
DRC: DNA replication checkpoint
HU: Hydroxyurea
RFB: Replication fork barrier
RS: Replication stress

## Material and Methods

### Yeast strains, plasmids and primers

All yeast strains in this study were generated in the W303 background (*ade2-1 ura3-1 his3-11, trp1-1 leu2-3, can1-100*) and are listed in Table 1.

**Table 1:**
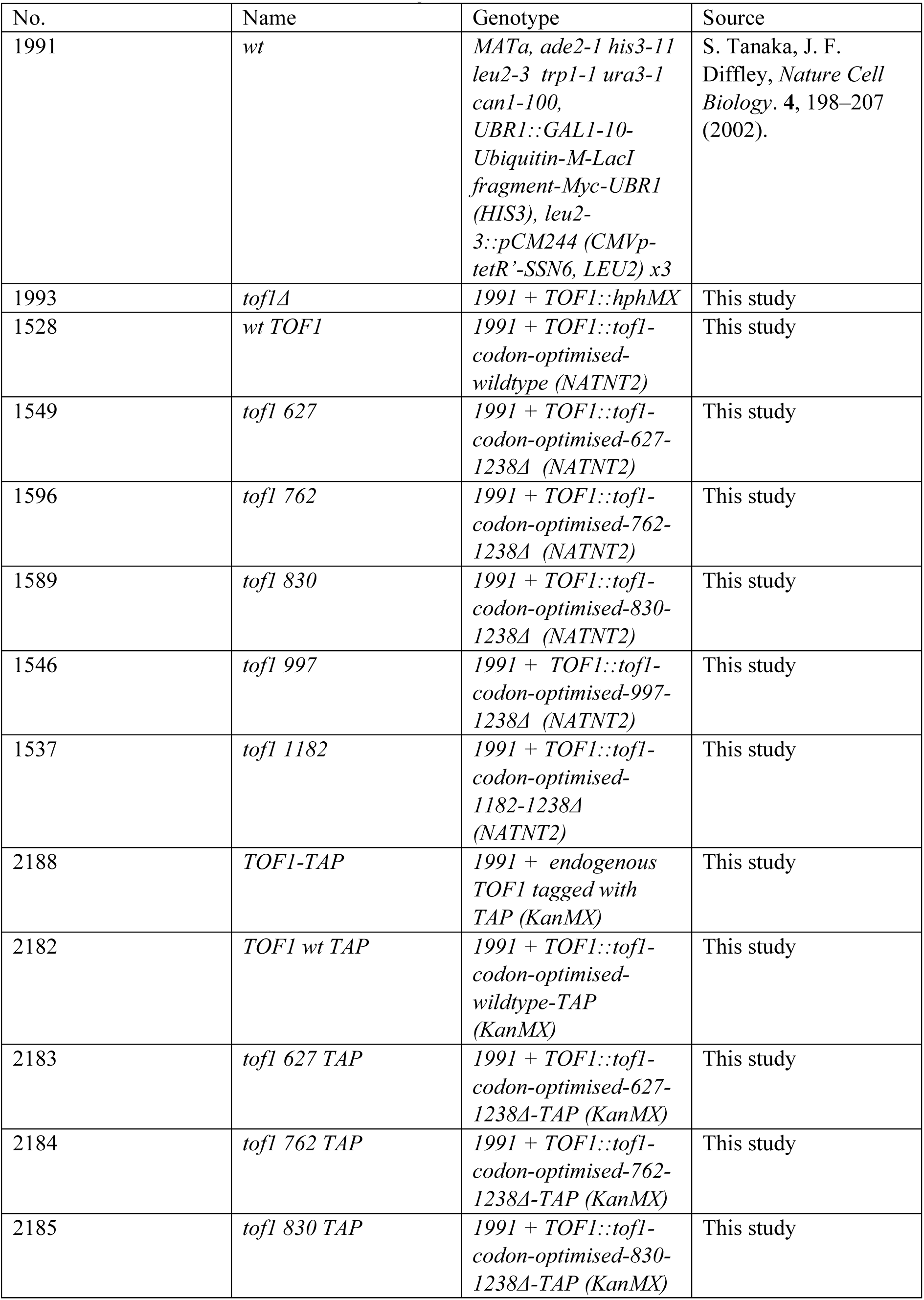
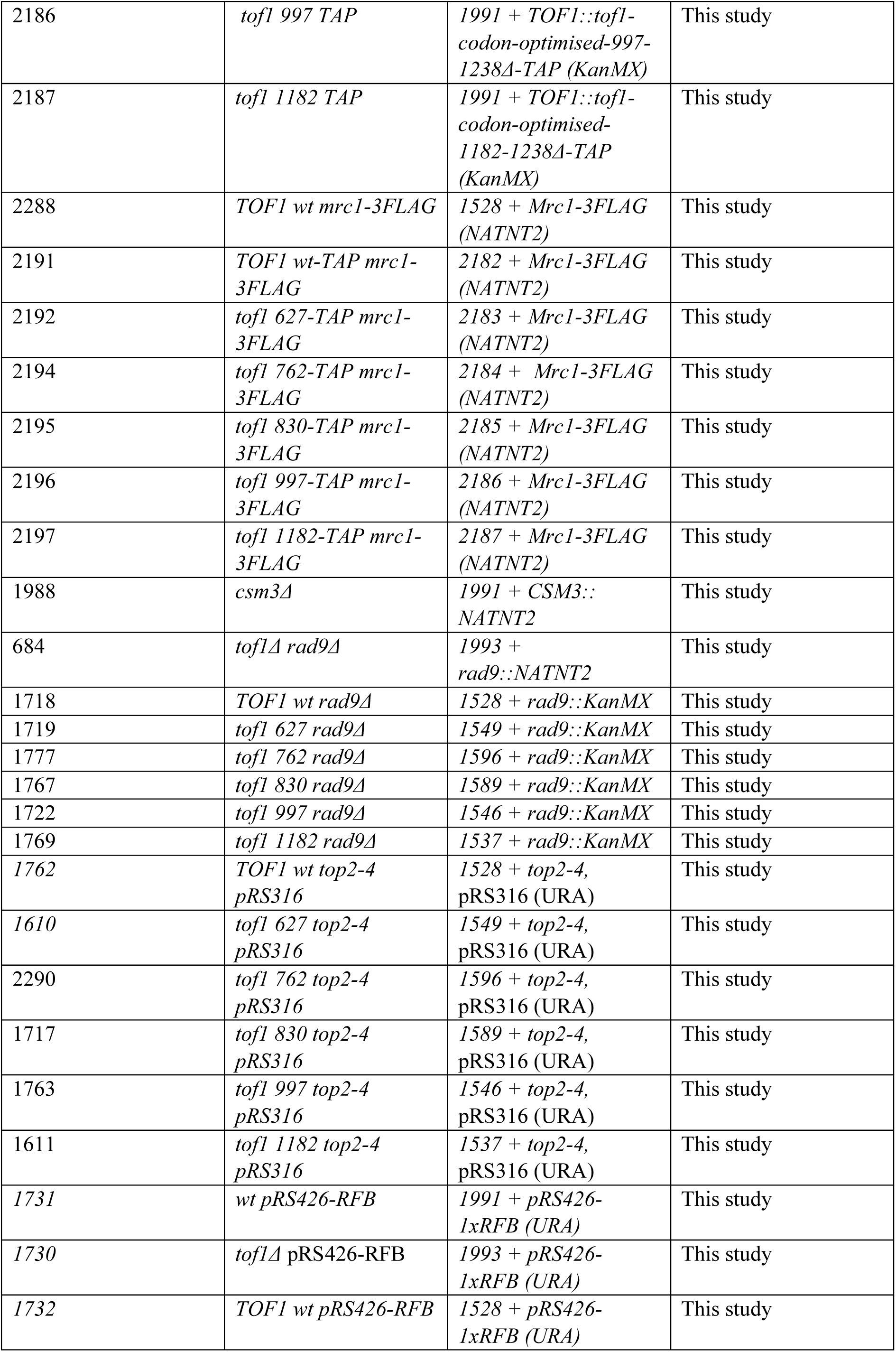
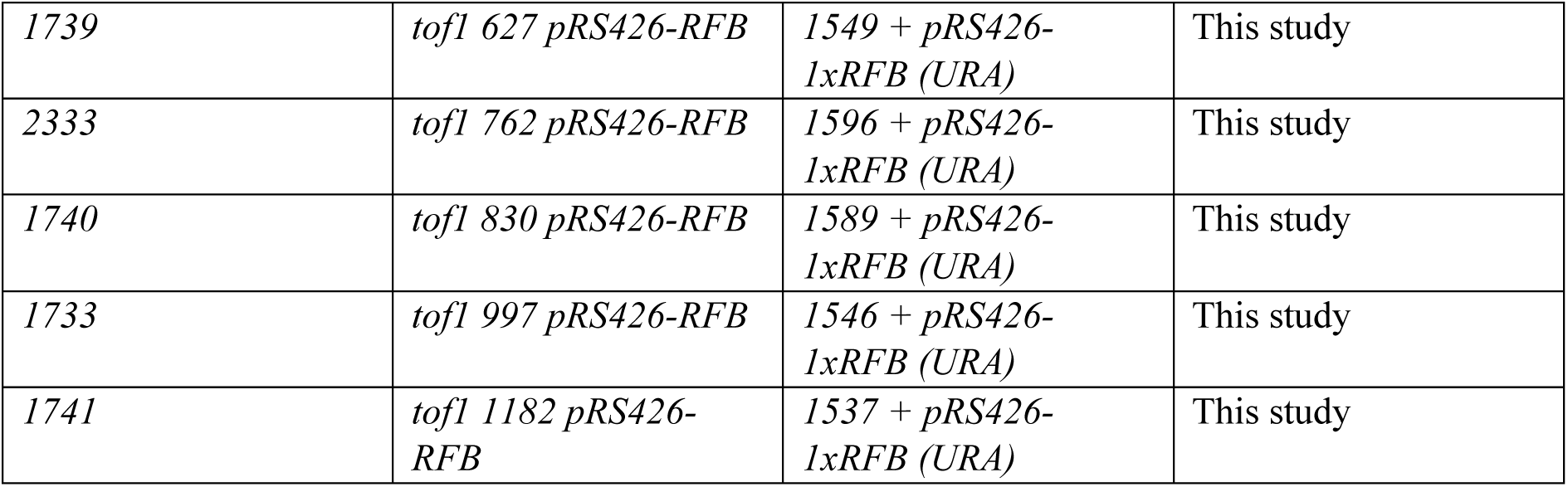
Yeast Strains Used in this Study.

See Table 2 for all plasmids used in this study.

**Table 2:** Plasmids used in this Study.

| Plasmid Name | Source | Used for |
| --- | --- | --- |
| pRS306/Tof1- Gal-CBP-Csm3 | Yeeles et al., 2017 | Generation of pBAK004 containing Tof1-codon optimised sequence upstream of NATNT2 selection marker |
| PBAK004 (TOF1-wt-NATNT2) | This study | Used as template for site-directed mutagenesis of TOF1 codon-optimised gene |
| tof1 627-NATNT2 | This study | Used as PCR template to generate fragment: tof1-codon-optimised-627-1238Δ-NATNT2 for yeast transformation into endogenous TOF1 locus |
| tof1 762-NATNT2 | This study | Used as PCR template to generate fragment: tof1-codon-optimised-762-1238Δ-NATNT2 for yeast transformation into endogenous TOF1 locus |
| tof1 830-NATNT2 | This study | Used as PCR template to generate fragment: tof1-codon-optimised-830-1238Δ-NATNT2 for yeast transformation into endogenous TOF1 locus |
| tof1 997-NATNT2 | This study | Used as PCR template to generate fragment: tof1-codon-optimised-997-1238Δ-NATNT2 for yeast transformation into endogenous TOF1 locus |
| tof1 1182-NATNT2 | This study | Used as PCR template to generate fragment: tof1-codon-optimised-1182-1238Δ-NATNT2 for yeast transformation into endogenous TOF1 locus |
| pRS316 | Sikorski, R. S. and Hieter, P. (1989) | Transformed into tof1 top2-4 strains for fork rotation/catenation assay |
| 4XRFB | Luis Aragon Lab (unpublished) | Used as PCR template to amplify 1xRFB sequence for cloning into pRS426 |
| pRS426-RFB | This study | Transformed into tof1 top2-4 strains for fork pausing assay |

Generation of yeast strains expressing truncated forms of Tof1 was carried out in two steps. First, codon-optimised *TOF1* gene was cut out from a pRS306/Tof1-Gal-CBP-Csm3 plasmid (gift from Diffley lab) (Yeeles *et al*, 2017) by NotI digestion and cloned into pFA6-natNT2. The codon-optimised TOF1 was then mutagenized using QuikChange Lightning Site Directed Mutagenesis kits (Agilent, 210518) according to manufacturer’s instructions, with primer sets designed to incorporate premature stop codons into the TOF1 open reading frame. Mutagenesis was confirmed by sequencing before PCR amplification and insertion of the mutagenized sequence and Nourseothricin resistance gene into the endogenous TOF1 locus of yeast cells using lithium acetate transformation.

Plasmid pRS426-RFB was generated by a 2-fragment Gibson assembly. Specifically, a single RFB site was PCR-amplified from plasmid 4xRFB (gift from Luis Aragon) corresponding to the sequence from *Saccharomyces cerevisiae* S288C Chromosome XII: 459799 - 460920. This was assembled with *PfoI*-linearized pRS426 using the NEBuilder® HiFi DNA Assembly Cloning Kit (New England Biolabs, E5520S).

### Media and Cell-Cycle Synchronisation

For plasmid catenation experiments in *top2-4* pRS316-containing strains, cells were grown to mid-log phase in synthetic complete media without uracil +2% glucose, before being re-suspended in in YP 2% Glucose (YPD). Cells were arrested in G1 by addition of 10 µg/ml alpha factor peptide (Genscript) for 1.5 hrs, after which a second dose of alpha factor (5µg/ml) was added. When >90% of cells were unbudded, cultures were shifted to the restrictive temperature for *top2-4* (37⁰C), for one hour before release into S-phase by washing three times with YPD at 37⁰C. Time 0 indicates the time from addition of the first wash. 50 µg/ml nocodazole was added to cultures at 45 minutes to prevent mitotic entry, and at 80 minutes from release 10 ml samples for 2D gel and Southern blotting analysis were collected by centrifugation and snap-freezing the pellets in liquid nitrogen.

For fork pausing experiments cells containing pRS426-RFB were grown to mid-log phase in synthetic complete media without uracil +2% glucose. 10ml cultures were collected by centrifugation and snap-freezing the pellets in liquid nitrogen.

For experiments involving treatment with HU, cells were grown in YPD to mid-log phase before either HU being added to 200 mM for 120 minutes (for Rad53 activation experiments in figure 5A) or for experiments started with a G1 arrest, mid-log cells were treated with 10 µg/ml alpha factor peptide. To release cultures into the cell cycle, when cells were >90% unbudded, cultures were washed 3 times in YPD and re-suspended after the third wash in YPD containing 200 mM HU. Time 0 was designated as the time from the first wash. Time points were taken as indicated. For release from HU experiments, HU-containing media was washed off of cells by washing 3 times with YPD before re-suspending in YPD. Time 0 is taken as the time from the first YPD wash.

For RPA ChIP experiments cells were grown in YP +2% raffinose to mid-log phase at 25 °C, before being arrested with 10 µg/ml alpha factor peptide. After 1hr 45 min 2% galactose and an additional 5 µg/ml alpha factor was added. After 2hr, when cells were >90% unbudded, 25 µg/ml doxycycline was added. 15 minutes after doxycycline addition temperature was switched to 37°C and incubated for 1hr. Cells were then released by washing 3 times with pre-heated YP 2% raffinose 2% galactose with 25 µg/ml doxycycline and resuspended in the same media supplemented with 200 mM HU. Time 0 was taken as the time from the first wash. Samples were then incubated for 1 hr at 37°C before being fixed by resuspending in YP + 1% formaldehyde (Sigma) for 45 min at 25 °C. 125 mM glycine was then added followed by a 5 min incubation at 25 °C. Cells were washed with PBS before being pelleted and snap-frozen in liquid nitrogen.

### TCA Extraction

10 ml of mid-log yeast cultures were centrifuged at 3500 rpm for 5 min and the resulting cell pellets were snap-frozen in liquid nitrogen before storing at −80°C. All further steps were carried out on ice, and centrifugation and homogenisation steps at 4°C. 200 µl of 20% TCA was added to thawed cell pellets and the cell suspension was transferred to screw-cap tubes containing 500 µl of 0.5mm zirconia/silica beads (BioSpec Products). Cells were homogenised using a FastPrep-24 (MP Biomedicals) on max speed (6.5 m/s) for 4 x 1 min pulses, with 1 minute on ice in between pulses. Beads were separated from the lysate by piercing the tubes and centrifugation of the mixtures into fresh tubes at 3000 rpm for 2 min. Beads were washed once with 600 µl 5% TCA and centrifuged again into tubes containing the cell lysate. Cell extract/TCA mixtures were then centrifuged at 13000 rpm for 5 min before removing all TCA from the resulting pellets. This step was repeated once to ensure complete removal of all TCA. To the pellets 200 µl of 1 X sample buffer was added before boiling samples for 5 min. Samples were spun at 13000 rpm for 5 min and the resulting supernatants were collected and stored at −20°C for SDS-PAGE and western blotting analysis.

### SDS-PAGE and western blotting

Protein extracts prepared by TCA extraction were run on either 8%, 10% or 12% SDS-PAGE gels before being wet-transferred to nitrocellulose membranes at 50V for 90 min, 4°C. Proteins were visualised by staining in Ponceau for 30 seconds before membranes were blocked in 5% milk (Marvel) PBS 0.2% Tween-20 (PBS-T). All primary antibodies were diluted in 5% milk PBS-T and incubated overnight at 4°C. In between primary and secondary antibody incubations membranes were washed 3 times with PBS-T for 15 min. Secondary antibodies were diluted in 5% milk PBS-T and incubated with membranes for 1 hr at room temperature. Proteins were detected using Western Lightning Plus-ECL (Perkin-Elmer, NEL104001EA) and images were acquired on an ImageQuant LAS4000 system (GE Healthcare). Densitometry analysis was carried out using ImageQuant TL software.

Primary antibodies used for immunoblotting in this study: OctA-Probe Antibody 1:1000 (for detecting FLAG-tagged proteins) (sc-166355), anti-Csm3 (Gambus *et al*, 2006), anti-PAP 1:1000 (for detecting TAP-tagged proteins) (Sigma-Aldrich, P1291), anti-Rad53 1:2000 (abcam ab104232). Secondary antibodies used in this study: anti-mouse-HRP 1:1000 (Dako P0260), anti-rabbit-HRP 1:1000 (Dako P0448) and anti-sheep-HRP 1:10000 (Sigma-Aldrich A3415).

### TAP pulldowns

200 ml of mid-log yeast cultures were centrifuged for 5 min at 3500 rpm and washed once in PBS. The resulting cell pellet was snap-frozen in liquid nitrogen and stored at −80°C before proceeding. Frozen pellets were re-suspended in 0.5 ml of ice-cold lysis buffer (50 mM HEPES pH 7.5, 300 mM KCl, 5 mM EDTA, 10% Glycerol, 80 mM Beta-glycerophosphate, 0.01% Triton X-100, 1 protease inhibitor tablet (cOmplete™, EDTA-free Protease Inhibitor Cocktail, Roche) and 1 phosphatase inhibitor tablet (PhosSTOP™, Roche)) in screw-cap tubes. 300 µl of zirconia/silica beads (BioSpec Products) were added and the cell suspensions were homogenised using the FastPrep-24 (MP Biomedicals) on max speed (6.5 m/s) for 4 pulses of 1 min each, with 3 min on ice in between each pulse. The resulting lysate was transferred to fresh tubes and cleared by centrifugation at 13000 rpm for 5 min. Cleared supernatants were then transferred to fresh tubes before starting the pulldown. 10 µl of IgG Sepharose 6 Fast Flow affinity resin (GE-Healthcare, #17096901) for each pulldown were pre-washed 3 times in cold lysis buffer before being added to the cleared cell lysates. The lysate/bead mixtures were incubated at 4°C on a rotating platform for 2 hrs and subsequently poured into Poly-Prep® Chromatography Columns (Bio-Rad #7311550). Unbound fractions were collected through the columns and stored at −20°C. The beads were washed 4 times in the columns with 1 ml of cold lysis buffer before the bound fraction was eluted in 0.5 ml of 0.2 M glycine pH 3.0. Input, unbound and eluted proteins were precipitated by TCA extraction before running on 4–20% Mini-PROTEAN® TGX™ Precast Protein Gels (Bio-Rad #4561096) and western blotting was performed as described.

### DNA Purification for Southern Blotting

Frozen pellets were re-suspended in 0.4 ml of DNA Extraction Buffer (50 mM Tris-HCl pH 8.0, 100 mM NaCl, 10 mM EDTA, 1% SDS) along with 40 Units of lyticase (Sigma-Aldrich, L2524) and 5 µl 2-mercaptoethanol (Sigma Aldrich, 63689). Samples were incubated at 37°C for 5 min before addition of 450 µl phenol/chloroform/isoamylalcohol (25:25:1, Sigma-Aldrich) and mixing by rotation. Phase lock tubes (5 Prime, 2302800) were used to collect the aqueous layer, by centrifugation for 5 min at 12000 rpm. DNA was ethanol precipitated by addition of roughly 2 x volume of 100% EtOH and washed once in 70% EtOH before air-drying and solubilisation in 10 mM Tris pH 8.0.

For plasmid DNA catenation analysis of pRS316, purified DNA was nicked with Nb.Bsm1 (New England Biolabs, R0706S) according to manufacturer’s instructions.

For fork pausing analysis of pRS416-RFB, purified DNA was digested with BamHI-HF (New England Biolabs, R3136S) and SnaBI (New England Biolabs, R0130L) according to manufacturer’s instructions.

After nicking/digestion the DNA was precipitated, washed and solubilised as above with the addition of 300 mM Sodium Acetate pH 5.2 at the first ethanol addition.

### 2-D Gel Electrophoresis of replication intermediates or catenated plasmid replication products

Purified DNA was separated in the first dimension by electrophoresis in 0.4% MegaSieve/MegaBase Agarose (Scientific Laboratory Supplies, H15608), 1X TBE (90 mM Tris, 90 mM Boric Acid, 10 mM EDTA).

For DNA catenation analysis of plasmid pRS316 from *top2-4* cells, 1^st^ dimension gels were run at room temperature for 16-18 hrs at 30V. A portion of the gel containing small amount of each DNA sample was excised and stained in 0.5 µg/ml Ethidium Bromide 1XTBE to reveal extent of genomic DNA mobility. The remaining non-stained gel slices containing the plasmid were excised and embedded in 1.2% MegaSieve/MegaBase Agarose 1X TBE and run in 1XTBE at 4°C for 16-17 hrs, 120V.

For analysis of paused replication intermediates from plasmid pRS426-RFB, for 1^st^ dimension gels were run at room temperature for 15.5 hrs at 30V. 1^st^ dimension gels were stained in 0.5 µg/ml Ethidium Bromide in 1XTBE and gel slices containing the replication intermediates were excised and embedded in 1% MegaSieve/MegaBase Agarose 1X TBE 0.3 µg/ml Ethidium bromide gels. 2^nd^ dimension gels were run in 1XTBE at 4°C for 8 hrs, 120V with re-circulation of the running buffer.

### Southern Blotting

Following 2-D electrophoresis, gels were washed sequentially in Depurination buffer (0.125M HCl), Denaturation buffer (0.5 M NaOH, 1.5 M NaCl) and Neutralisation buffer (0.5 M Tris-HCl, 1.5 M NaCl pH 7.5) with washes in ddH_2_O in-between each buffer. DNA was transferred onto Hybond-N+ membrane (GE Healthcare) by capillary action in 20X SCC (3M NaCl, 350 mM NaOC Trisodium Citrate pH 7.0). Membranes were cross-linked using a UV Stratalinker 1800 (Stratagene) at 1200J/m and subsequently blocked in hybridisation buffer (5X SSC, 5% Dextran sulphate (Sigma-Aldrich, D8906) 0.2% Tropix I-Block (Applied Biosystems, T2015), 0.1% SDS) for at least 1 hr at 60°C.

Catenated pRS316 plasmids or replication intermediates from pRS416-RFB were probed with DNA amplified from pRS316 (probing specifically for the URA3 gene). Labelling and detection used random prime labelling incorporating fluorescein tagged dUTP (Roche). Following probing, hybridized fluorescein tagged dUTP was detected with alkaline phosphatase tagged anti fluorescein Fab fragments (Roche), revealed with CDP-Star (GE Healthcare) and non-saturating exposures acquired on an ImageQuant LAS4000 system (GE Healthcare). Densitometry analysis was carried out using ImageQuant TL software.

### FACS analysis of DNA content

For analysis of cell cycle progression, 0.5 ml of yeast culture was pelleted by centrifugation at 13000 rpm for 15 seconds before removal of all growth media. The pellets were re-suspended in 0.5 ml 70% ethanol and stored at 4°C before processing and analysis.

Fixed cells were washed in 50 mM Tris pH 8.0 and 10 mg/ml RNaseA (Sigma-Aldrich) was added. Cells were incubated overnight at 37⁰C, pelleted and re-suspended in freshly made 5 mg/ml pepsin in 5 mM HCl and incubated again at 37⁰C for 30 min. Fixed cells were washed once more in 50 mM Tris pH 8.0 and re-suspended in 0.5 mg/ml propidium iodide in 50mM Tris pH 8.0. Samples were sonicated for 5 seconds each on low power to reduce clumping before analysis using the BD Accuri™ C6 Plus (BD Biosciences).

### Drug sensitivity assays

Yeast cells were grown to mid-log phase before being serially diluted 10-fold in YPD. 5 µl of each dilution was spotted onto YPD plates containing the indicated dose of drug or control reagent and incubated for 24-28 hr at 25°C before imaging.

### Colony Survival Assays

Yeast cells were grown to mid-log phase before being arrested in G1 by the addition of 10 µg/ml alpha factor peptide. When cells were >90% unbudded they were released into the cell cycle in the presence of 200 mM HU for 1 hr. Following the HU treatment cells were counted, diluted in YPD medium and plated onto YPD plates. Colonies were counted 48 hrs after plating and the viability was calculated as the percentage of plated cells able to form colonies. Statistical significance was calculated using an unpaired Students t-test.

### RPA1 ChIP-seq

Pellets from 75 ml cultures were resuspended in 500 μl SDS buffer (1% SDS, 10 mM EDTA, 5M Tris HCl, cOmplete Tablets Mini EDTA-free EASYpack (Roche), PhosSTOP (Roche)). 200 μl of 0.5 mm zirconia/silica beads were added to samples and cells were lysed using the FastPrep-24 (MP Biomedicals) on max speed (6.5 m/s), with 5 rounds of 1 min each. Lysate was spun out and IP buffer (0.1% SDS, 1.1% Triton-X-100, 1.2 mM EDTA, 16.7 mM TRIS HCl (pH8), cOmplete Tablets Mini EDTA-free EASYpack (Roche), PhosSTOP (Roche)) was added to a final volume of 1 ml. Samples were sonicated using the Focused-Ultrasonicator (Covaris) (Average incident power – 7.5 Watts, Peak Incident Power – 75 Watts, Duty Factor – 10 %, Cycles/Burst – 200, Duration – 20 min). The sample was centrifuged for 20 min at 13000 rpm at 4°C. Supernatant was then diluted to 7.5 ml with IP buffer. 75 μl protein A Dynabeads (Invitrogen) and 75 μl protein G Dynabeads (Invitrogen), were washed 3 times in IP buffer before adding to the sample and incubating for 2 h at 4°C. 2 ml of the supernatant was taken to 15 ml falcon tubes, and the rest was kept at −20°C as an input sample. To the 2 ml sample RPA1 antibody (1:10000, Agrisera, AS07214) was added followed by overnight incubation on a rotating wheel at 4 °C.

A mix of Dynabeads, Protein A (30 μl) and Protein G (30 μl), was washed 3 times in IP buffer. This was added to each sample and incubated at 4°C for 4 h. Supernatant was removed and beads were washed at 4°C for 6 min in TSE-150 (1% Triton-X-100, 0.1% SDS, 2 mM EDTA, 20 mM Tris HCl (pH8), 150 mM NaCl), followed by TSE-500 (1% Triton-X-100, 0.1% SDS, 2 mM EDTA, 20 mM Tris HCl (pH8), 500 mM NaCl), followed by LiCl wash (0.25 M LiCl, 1% NP-40, 1% dioxycholate, 1 mM EDTA, 10 mM Tris HCl (pH8)) and finally Tris-EDTA (TE pH8). Elution was carried out in 400 μl elution buffer (1% SDS, 0.1M NaHCO3, for 30 min at room temperature. At the same time 50 μl from the input sample was added to 150 μl of elution buffer. 20 μl of 5 M NaCl and 10 μl of 10 mg/ml proteinase K (Invitrogen) was then added to the input, and 40 μl and 20 μl to the IP samples respectively. These were incubated at 65°C overnight. Then 10 μl of DNase-free RNase (Roche) was added to the input and 20 μl to the IP samples, and they were left at 37°C for 30 min. All DNA was purified with a Qiagen PCR purification kit and eluted in 40 μl H_2_O. 34 ul from the RPA1 samples and 1 ng DNA in 34 ul water from the input were used for library preparation. 5 µl 10 x NEB2.1 buffer and 5 µl of random primers (8N, 3 mg/ml stock) were added and the samples were boiled at 95°C for 5 min and immediately placed on ice for 5 min. 5 µl 10 x dNTPs with dUTP instead of dTTP (2 mM each) and 1 µl T4 polymerase (NEB) were added and the mixture was incubated at 37°C in a thermal cycler for 20 min, and 5 µl 0.5 M EDTA (pH 8) was immediately added to stop the reaction. The resulting dsDNA was used to create libraries using the Ultra II library kit (NEB) as per the manufacturer’s instructions with 13 cycles at the amplification step.

Paired end sequencing was performed using the MySeq (75bp reads from each side) or NextSeq 500 (42 bp reads from each side) systems to result >2 million reads.

### ChIP-SEQ analysis

FASTQ files were generated by Illumina basespace (https://basespace.illumina.com/home/index). The resulting sequences were aligned to a reference genome (R64-1-1, *Saccharomyces cerevisiae* S288c assembly from Saccharomyces Genome Database) using Bowtie 2 generating a SAM output file for each sample (http://bowtie-bio.sourceforge.net/bowtie2/index.shtml). Reads from MySeq were trimmed 25 bp from 3’ and 1 bp from the 5’ end, while reads from NextSeq were not trimmed.

Command for MySeq reads:

~~~
  bowtie2 -p 14 -x [path to index folder] --trim3 25 --trim5
1 −1 [Path and name of R1 fastq file] −2 [Path and name of R2
fastq file] -S [name of the resulting .sam file]
~~~

Command for NextSeq reads:

~~~
  bowtie2 -p 14 -x [path to index folder] --trim3 0 --trim5
0 −1 [Path and name of R1 fastq file] −2 [Path and name of R2
fastq file] -S [name of the resulting .sam file]
~~~

SAM files were then converted into sorted BAM files by using SAMtools (http://samtools.sourceforge.net/):

~~~
  samtools sort [name of the .sam file generated with
bowtie2] -o [name for the resulting .bam file] -O bam -T [name
for resulting .bam file wo .bam]
~~~

Duplicates were then removed using picard (https://broadinstitute.github.io/picard)

~~~
java -jar ∼/picard/picard-tools-1.138/picard.jar
MarkDuplicates I= [name for the resulting .bam file]    O=
[name for the resulting without repeats.bam file] M= [name of
metrix file.txt] REMOVE_DUPLICATES=true
~~~

BAM files were used for Model-based Analysis of ChIP-Seq (MACS2). We used the ‘call peak’ function which also generates genome wide score data. These were used to generate fold enrichment tracks. Example command:

~~~
  macs2 callpeak -t [sorted BAM file from yh2a data]-c
[sorted BAM file from h2a data]-f BAMPE -g 12100000 -n [name
for output file] -B -q 0.01 --SPMR
~~~

The data then was sorted into 50 bp bins, normalized to have a mean value of 1, and used for meta data analysis using custom-made R programs.

### Sync-SEQ

Pellets from 2 ml cultures were resuspended in 500 μl SDS buffer (1% SDS, 10 mM EDTA, 5M Tris HCl, cOmplete Tablets Mini EDTA-free EASYpack (Roche), PhosSTOP (Roche)). 200 μl of 0.5 mm zirconia/silica beads were added to samples and cells were lysed using the FastPrep-24 (MP Biomedicals) on max speed (6.5 m/s), with 5 rounds of 1 min each. Lysate was spun out and IP buffer (0.1% SDS, 1.1% Triton-X-100, 1.2 mM EDTA, 16.7 mM TRIS HCl (pH8), cOmplete Tablets Mini EDTA-free EASYpack (Roche), PhosSTOP (Roche)) was added to a final volume of 1 ml. Samples were sonicated using the Focused-Ultrasonicator (Covaris) (Average incident power – 7.5 Watts, Peak Incident Power – 75 Watts, Duty Factor – 10 %, Cycles/Burst – 200, Duration – 20 min for G1 samples and 13.5 for HU and released samples). 200 ul of sample was taken out and 20 μl of 5 M NaCl and 10 μl of 10 mg/ml proteinase K (Invitrogen) was then added followed by an overnight incubation on 65°C. Then 10 μl of DNase-free RNase was added to the sample and they were incubated at 37°C for 30 min. DNA was then purified with a Qiagen PCR purification kit and eluted in 50 μl H_2_O. 50 ng of DNA in 50 ul water was used for library preparation using the Ultra II library kit (NEB) as per the manufacturer’s instructions with 6 cycles at the amplification step.

Paired end sequencing was performed using NextSeq 500 (42 bp reads from each side) systems to result >2 million reads.

### Sync-SEQ analysis

Sync-seq analysis was done by using LocalMapper shell script and Repliscope R package from https://github.com/DNAReplicationLab/ (Dzmitry G Batrakou^1^, Carolin A Müller^1^, Rosemary H C Wilson^1^ and Conrad A Nieduszynski, DNA copy number measurement of genome replication dynamics by high-throughput sequencing – the sort-seq, sync-seq and MFA-seq family – accepted to Nature protocols)

~~~
localMapper.sh -g [path to index folder] [Path and name of R1
fastq file] −2 [Path and name of R2 fastq file] -s [name of
the output files] -w 3000 -c 14
~~~

The resulting .bed files were then read in to R:

~~~
repBed <- loadBed(file name for the replicating sample)
nrepBed <- loadBed(file name for the non-replicating sample)
~~~

Outliers were removed:

~~~
repBed<-rmOutliers(repBed, “median”, loLim = 0.25)
repBed<-rmOutliers(repBed, “max”, n = 2)
~~~

~~~
nrepBed<-rmOutliers(nrepBed, “median”, loLim = 0.25)
nrepBed<-rmOutliers(nrepBed, “max”, n = 2)
~~~

Ratio between non-replicated and replicated samples were calculated:

~~~
ratio <- makeRatio(repBed, nrepBed)
~~~

And normalised:

~~~
ratio <- normaliseRatio(ratio, [rFactor])
~~~

Where rFactor was empirically determined to fit the lowest replicating regions to 1.

The resulting ratios were smoothed by a moving average of 2 and plotted using custom-made R programs.

## Acknowledgments

We thank John Diffley for TOF1 coding sequence, Luis Aragon for budding yeast RFB DNA sequence and Karim Labib for antibodies to Csm3. We thank Maksym Shyian David Shore, Conrad Nieduszynski and Joe Yeeles for communicating data prior to publication. We thank Bonnie Brewer for advice on interpreting 2D agarose gels. This work was funded by the Biotechnology and Biological Sciences Research Council, United Kingdom, (BBSRC UK) Grants ref BB/J018554/1 (A.K., N.M., J.B.) and by a generous donation from Kate Fickle and Jerry Carroll (American Friends of University of Sussex) (R.W.).

## Author Contributions

R.W. and D.J. generated all the TOF1 truncation mutations. R.W. performed all pull downs and western analysis, 2D agarose gel electrophoresis and Southern blotting for replication pauses and DNA catenation, viability spot tests and Rad53 activation experiments. A.K. and N.E.M performed Sync-SEQ and ChIP-SEQ experiments. J.B. conceived and co-ordinated the study. J.B. wrote the manuscript with all authors contributing.

## Data Availability

All strains are available on request. All sequencing is available on request.

